# TEscape: Defining the human transposable element transcriptome using multiplatform long-read sequencing

**DOI:** 10.64898/2026.07.08.737305

**Authors:** Rafael L. V. Mercuri, Daniela M. Mombach, Felipe R. C. dos Santos, Joaquín Pérez-Schindler, Yi Huang, Pieter Spealman, Greta Pintacuda, Aziz Al’Khafaji, Elisa R. Donnard, Melina Claussnitzer, Pedro A. F. Galante

## Abstract

Transposable elements (TEs) not only account for half of the human genome sequence but also generate transcripts that contribute to transcriptomic diversity. Yet, their repetitive nature has hindered accurate quantification of the full TE-derived transcriptome, a challenge that long-read sequencing can overcome. Here, we combined multiplexed arrays isoform sequencing (MAS-ISO-seq) with a dedicated computational framework (TEscape) to perform an in-depth annotation of the human TE transcriptome. To capture the breadth of human transcriptome diversity, we profiled six representative cell types spanning three distinct biological contexts, including metabolism with, primary patient-derived adipogenic cells at two differentiation stages, and iPSC derived hepatic progenitor cells; the nervous system with iPSC-derived neurons, neural progenitor cells (NPCs), and pluripotency using induced pluripotent stem cells (iPSCs). Together, these datasets yielded over 235 million full-length long reads. First, to assess data coverage and transcriptome depth, we quantified protein-coding gene expression, detecting 14,312 genes (73.6% of all annotated protein-coding genes), which is a level consistent with deep and comprehensive transcriptome representation. Second, focusing on TE-derived transcripts, we identified >83,000 previously unannotated isoforms, the vast majority (84%) originating from a complex combination of multi-TEs. We also identified solo TEs, which are predominantly from LINE1 (14%). We confirmed that TE-transcripts are able to be exemplified by signatures detected in Liver Hepatocellular Carcinoma (LICH). Together, MAS-ISO-seq and TEscape establish the first long-read-based, high-resolution atlas of transcribed human TEs, providing a foundational resource for integrative transcriptome analyses and for investigating TE expression and regulation in health and disease.

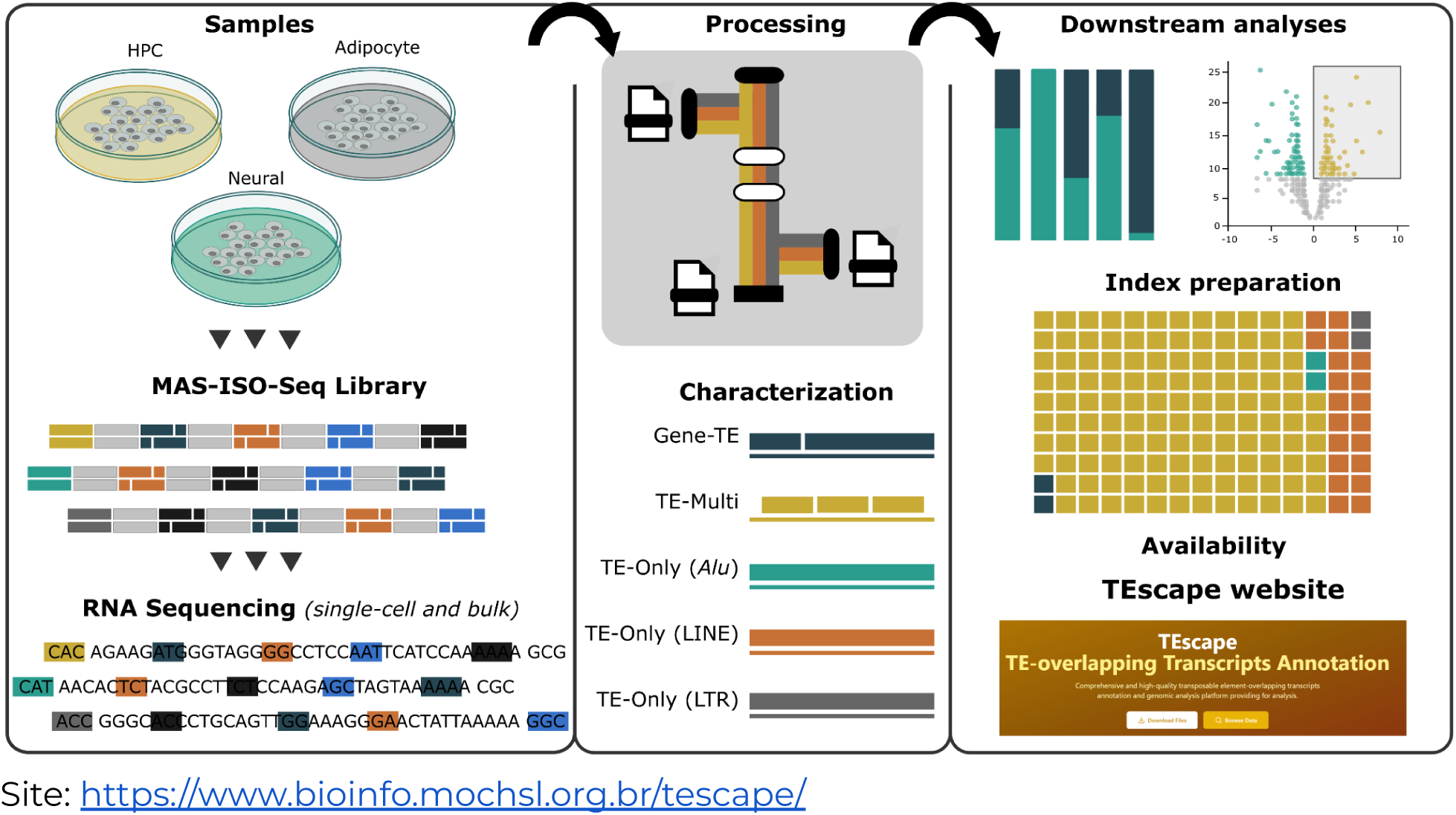
Graphical Abstract.

## INTRODUCTION

The human genome harbors a remarkable complexity that extends far beyond its protein-coding genes. While protein-coding sequences constitute merely 1.3% of the genome, the vast majority of our DNA consists of non-coding elements, with transposable elements (TEs) comprising approximately 50% of its total (Lander et al. 2001). These mobile genetic elements, first discovered in maize (McCLINTOCK 1950), have traditionally been viewed as “junk DNA” or genomic parasites. However, recent advances in genomic technologies have revolutionized our understanding of their activities and functional significance in many species, including the *Homo sapiens*.

In humans, the most abundant TEs include Long and Short Interspersed Nuclear Elements (LINEs and SINEs, respectively), Long Terminal Repeat (LTR) retrotransposons and DNA transposons. While most of these elements are now inactive due to accumulated mutations, mounting evidence suggests they play crucial roles in genome organization, contribution to species evolution, gene expression regulation, and disease association (Mills et al. 2007; Brouha et al. 2003). Specifically, the use of Next-Generation Sequencing in transcriptomics (Wang et al. 2009) has revealed that TEs are not transcriptionally inert but contribute significantly to the human transcriptome diversity (Solovyov et al. 2025). TEs can be transcribed as standalone units or as part of larger transcripts, potentially generating novel (chimeric) mRNA with functional implications (Li et al. 2025; Panda and Slotkin 2020). However, the precise characterization of these TE-derived transcripts has been hampered by technical limitations inherent to short-read sequencing technologies, particularly in resolving repetitive sequences and determining their exact genomic origins (Li et al. 2025). Currently, a comprehensive catalog of TE-derived transcripts in the human transcriptome still remains elusive. Understanding the full extent of TE transcription and their complete sequences seems to be crucial for several reasons: (1) TE-derived transcripts may have regulatory functions in gene expression in physiological and development contexts; 2) they might contribute to cellular diversity across tissues; and 3) their expression or dysregulation may have implications in pathology (Gebrie 2023; Mika and Lynch 2022; Tam et al. 2019).

The emergence of long-read sequencing (LRS) has revolutionized not only genomic but transcriptomic research, by enabling unprecedented insights into complex genomic regions, such as repetitive ones. The ability to sequence full-length transcripts overcomes an intrinsic limitation of short-read RNA sequencing and can be used to assess the vast diversity of common and rare transcripts (Anvar et al. 2018; Woolley et al. 2025). However, traditional LRS is still relatively low-throughput (Hardwick et al. 2019), posing a cost barrier to a more thorough quantification of the human transcriptome. Recently, the development of Multiplexed Arrays Isoform Sequencing (MAS-ISO-Seq), provided an unprecedented opportunity to overcome this throughput limitation (Al’Khafaji et al. 2024). This technology combines massive multiplexing and deep coverage with full-length transcript resolution, enabling accurate quantification of complex and low-abundance transcripts.

In this study, we applied MAS-ISO-Seq across three major biological domains: pluripotency (iPSCs), metabolism (primary adipocyte progenitors, adipocytes, and iPSC-derived HPCs), and the nervous system (iPSC-derived neural progenitors and neurons). This approach encompassed both patient-derived primary cells and reprogrammed models, integrating single-cell and bulk transcriptomic profiling to comprehensively map the human repertoire of TE-derived transcripts (retrotranscriptome). We applied multiple analytical metrics, including comparisons with GENCODE annotations, integration of transcriptional marker databases, and validation using external short-read datasets to confirm our findings. Our results provide an in-depth view of the repertoire of TE-derived transcripts across distinct cell types and expand the known human transcriptome, uncovering additional complexity in biological processes and pathological mechanisms.

## RESULTS

### Comprehensive annotation of retroelements using MAS-ISO-Seq

To construct a comprehensive annotation of TE-derived transcripts, we leveraged MAS-ISO-Seq across six cell types. MAS-ISO-seq enables high-throughput, full-length RNA isoform sequencing (Al’Khafaji et al. 2024), making it ideally suited for our goal of defining the human TE transcriptome. MAS-ISO-seq can be applied to diverse RNA-seq modalities, and this technical diversity is reflected in our experimental design (**Figure 1a** and **Supplementary Table 1**).

**Figure 1.**
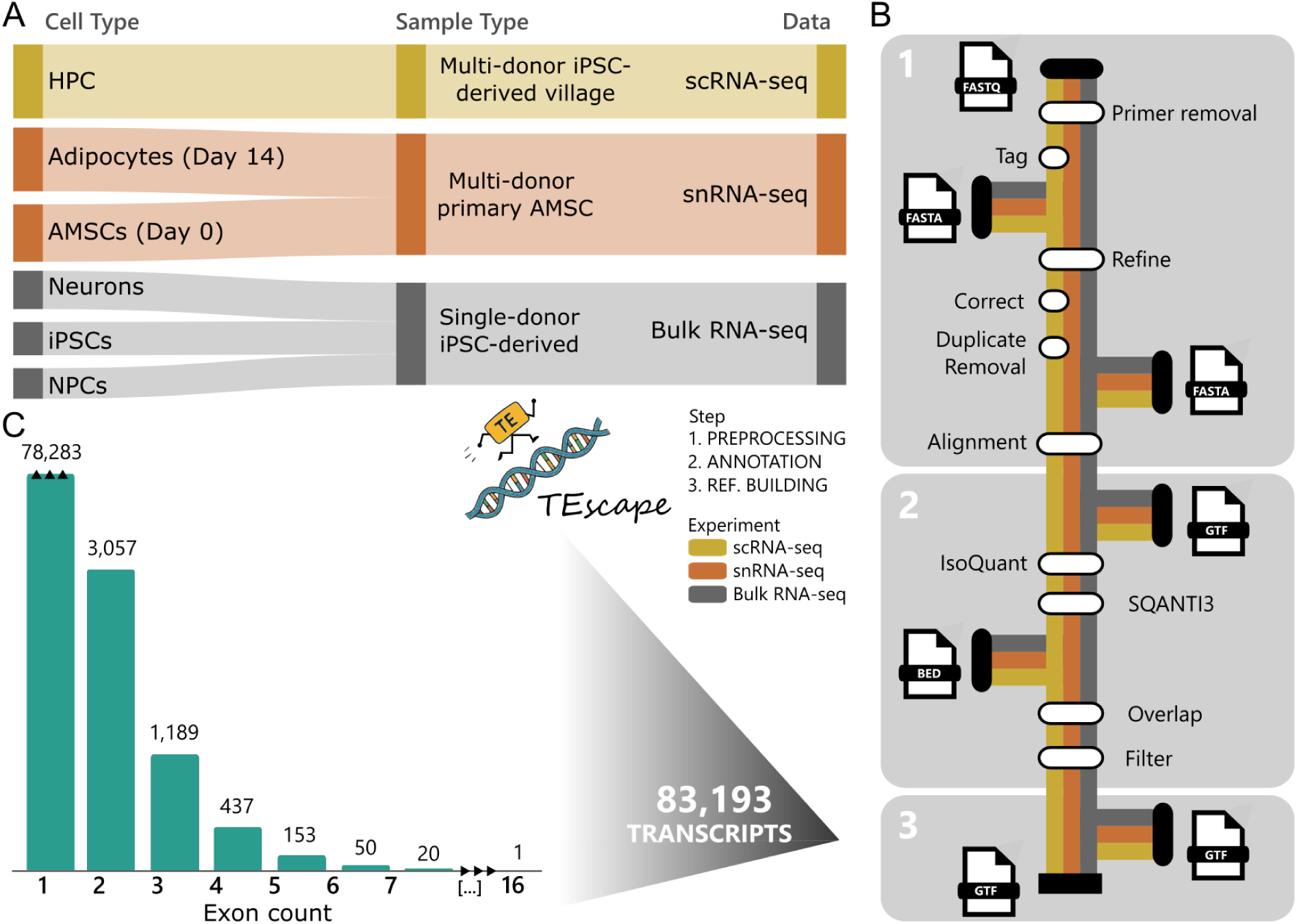
Overview of TEscape TE-derived transcripts annotation. **a)** MAS-ISO-seq samples across six cell types generated over 235 million long-reads. **b)** Pipeline steps: (i) preprocessing and alignment to hg38, (ii) transcriptome assembly using Gencode v47 and filtering for canonical junctions, followed by overlap analysis with RepeatMasker and RCPedia2.0, and (iii) integration of TE-derived transcripts into a new reference GTF. **c)** Final dataset includes 83,193 TE-derived transcripts, mostly single-exon isoforms. HPC = hepatic progenitor cells; iPSCs = induced pluripotent stem cells; NPCs = neural progenitor cells; scRNA-seq = single-cell RNA-sequencing; snRNA-seq = single-nucleus RNA-sequencing.

This methodological diversity was intentionally aligned with the cellular heterogeneity of the samples, allowing us to capture a broad spectrum of transcriptional phenotypes and maximize the detection of the variety of TE-derived transcripts. Specifically, we profiled hiPSC-derived hepatic progenitor cells (HPCs, day 7), primary adipocyte progenitors AMSCs (day 0) and differentiated adipocytes (day 14), induced pluripotent stem cells (hiPSCs) and two hiPSC-derived neural cell states: excitatory neurons (day 31) and neural progenitor cells (NPCs, day 4). HPCs were differentiated from a pool of hiPSCs from 89 donors using the Cell Village platform (Wells et al. 2023) and profiled via single-cell RNA-seq. Primary AMSCs were pooled from two donor samples, differentiated into adipocytes and profiled using single-nucleus RNA-seq (**Supplementary Figure 1**). NPCs and neurons were differentiated from a single hiPSC donor and profiled using bulk RNA-seq.

In total, we generated over 235 million long reads across all samples (**Supplementary Table 1**), establishing a robust foundation for the transcriptome-wide annotation of TE-derived RNAs. To systematically identify and characterize these transcripts, we implemented a three-step pipeline (**Figure 1b**). First, during the preprocessing phase, MAS-ISO-seq data were cleaned and refined using the Pigeon pipeline available at (https://github.com/PacificBiosciences/pigeon), including demultiplexing, barcode correction, and alignment to the hg38 reference genome. Second, in the annotation step, transcriptomes were assembled using IsoQuant (Prjibelski et al. 2023) with Gencode v47 (Mudge et al. 2025) as the reference, followed by stringent filtering with SQANTI3 (Pardo-Palacios et al. 2024) to retain only high-confidence isoforms, those with canonical splice junctions and no evidence of reverse transcription artifacts. Transcripts were then intersected with TE coordinates from RepeatMasker available at: www.repeatmasker.org, retaining only those with ≥60% overlap and supported by at least five reads. While several long-read analysis tools, such as FLAIR, commonly adopt a minimum threshold of three reads as independent observations of a given event, we applied a more stringent cutoff. Given our focus on monoexonic transcripts, which are more susceptible to spurious detection, we required a minimum of five supporting reads to increase confidence in transcript identification. In addition, for Village-derived data, transcripts were required to be present in at least two donors. Finally, in the reference-building step, we integrated the curated TE-derived transcripts into Gencode v36, v47 and v48 annotations and generated an updated GTF file.

This approach resulted in the identification of over 83,000 TE-derived transcripts (with median length 1192), the majority of which were single-exon isoforms but reaching up to 16 exons (**Figure 1c**), highlighting the extensive and previously underappreciated contribution of transposons to human transcriptome complexity.

Furthermore, to evaluate whether the sequencing depth was sufficient to capture the transcriptome complexity, we performed a saturation analysis (**Supplementary Figure 2**) across all samples. The results showed that isoform detection saturation approached 90% for all datasets, indicating comprehensive coverage.

### Retroelements improve gene annotation and exhibit transcription markers

To assess overlap with existing annotations, we compared our annotated transcripts against GENCODE v47, a comprehensive and up-to-date transcriptome reference, and RepeatMasker, the predominant annotation resource for mobile elements used by established in-locus TE quantification tools (TEtranscripts, Telescope, and SalmonTE).

We identified 1,258 transcripts previously annotated, corresponding to 773 genes present in GENCODE (**Figure 2a**). At the exon level, our annotation recovered approximately 6,690 exons already present in GENCODE. While these numbers represent only a small fraction when compared to the ∼83,000 transcripts and 87,891 exons annotated in TEscape, they highlight the consistency of our strategy with the growing evidence that transcriptome expansion is largely driven by TE-derived transcripts.

**Figure 2.**
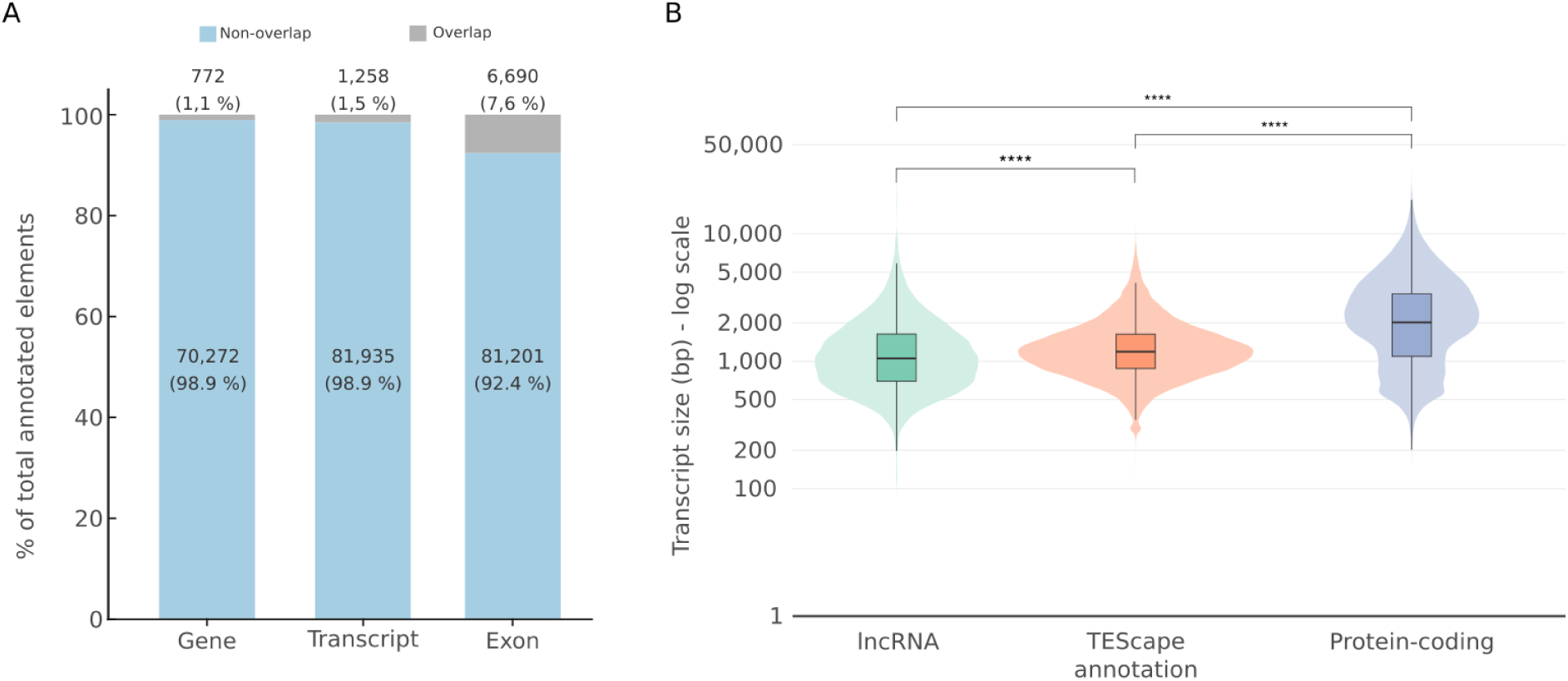
Gencode v47 transcripts comparison. **A**) Proportion of genes, transcripts, and exons showing overlap with transposable elements (TEs). Each bar represents the percentage of overlapping (dark blue) and non-overlapping (light blue) regions within each genomic feature. **B**) Transcript length distributions across biotype categories. Violin plots depict the size of transcripts annotated as long non-coding RNAs (lncRNAs), TE-derived transcripts (TEscape annotation), and protein-coding genes. Boxes indicate the interquartile range and median, while whiskers extend to 1.5× the interquartile range. Statistical significance between groups was assessed using the Wilcoxon rank-sum test (****p < 0.0001).

In addition, when comparing the average length of these transcripts and exons with those of lncRNAs and protein-coding genes also detected by our strategy, we observed statistically significant differences across the groups (**Figure 2b**). This suggests that TE-derived transcripts can be considered as a distinct class within the transcriptome. Notably, this pattern remained consistent even when restricting the comparison to transcripts already annotated in GENCODE, reinforcing the robustness of the observation (**Supplementary Figure 3**).

When compared against RepeatMasker, we required a minimum of 95% genomic overlap between annotated TE loci and our transcript models. Under this criterion, 666 TEScape transcripts exhibited near-complete concordance with annotated TE regions. This pattern suggests that these transcripts likely represent more complex TE-derived structures that are not adequately captured by conventional annotations, highlighting the need for more refined and curated transcript models.

We assessed whether the TE-derived transcripts exhibited regulatory expression features, including transcription start sites (TSS), promoters, distal or proximal enhancers, and CTCF binding sites, using data from the ENCODE (ENCODE Project Consortium et al. 2020) and TSS databases (Wakaguri et al. 2008). Remarkably, around 85% of transcripts in all datasets contained at least one of these markers (**Table 1**). We tested the randomness of this observation, by selecting a random set of similar mobile elements from RepeatMasker and comparing them with these same ENCODE and TSS data, we would observe a similar pattern. To test this, we performed a Monte Carlo simulation with 10,000 iterations, obtaining a p-value < 0.001. These findings show a widespread regulatory potential among TE-derived transcripts and indicate their expression relevance.

**Table 1.**
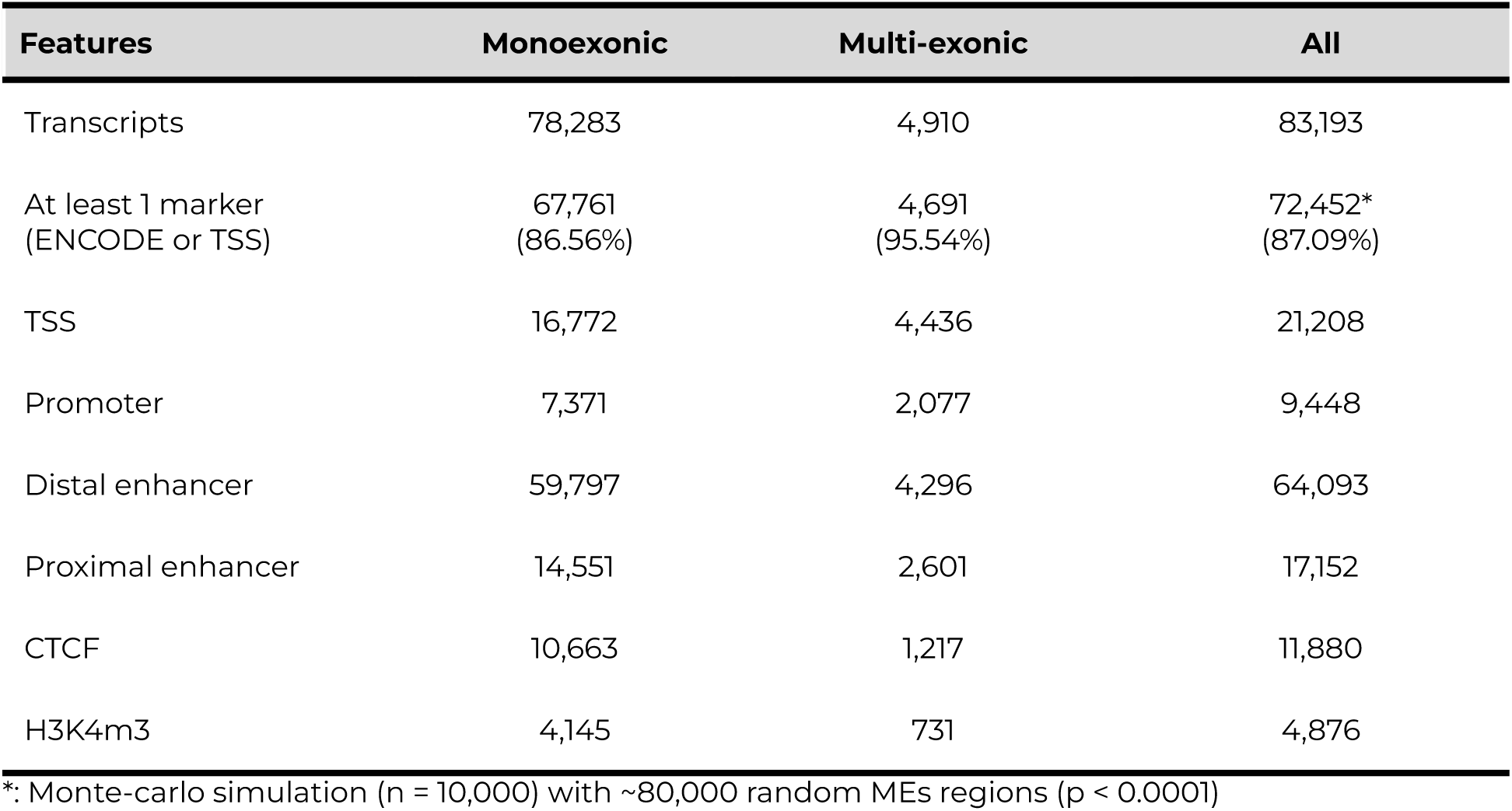
Expression markers (ENCODE and TSS database)

Notably, TE-derived transcripts were more accurately annotated using long-read sequencing, which provided enhanced resolution of transcription start and termination sites. This precision is critical given that TEs can hijack nearby regulatory elements at their insertion sites, leading to transcription initiation upstream of the TE sequence, sometimes extending several base pairs, as observed in *transcript101960.chr7.nnic* (L1PA3 transcript), and termination downstream, resulting in so-called “leaky expression” or 3′ run-on transcripts, such as transcript *transcript107879.chr5.nnic* (L1MA4 transcript), which ends 295 bp beyond the TE sequence boundary. Importantly, these transcripts had never been previously annotated, underscoring the value of long-read data followed by our strategy in refining transcript annotations (**Supplementary Figure 4**).

### TE-derived transcripts annotation exhibit both shared and cell type–specific patterns

Next, to investigate the cell type specificity of TE-derived transcripts, we examined transcript overlap across all six human cell types (**Figure 3**). Rather than a globally shared pattern, the intersection pattern captures the three biological contexts represented in this study. Within the metabolic lineage, AMSCs and differentiated adipocytes shared the largest pairwise intersection (27,258 transcripts), reflecting a conserved mesenchymal TE program across differentiation states. Critically, cross-cell-type sharing was highest within the metabolic lineage overall, i.e. HPCs, AMSCs and adipocytes shared more transcripts with each other than any metabolic cell type shared with cells of the neural lineage or iPSCs, indicating that TE transcriptional programs are organized along biological function rather than cell-of-origin. HPCs nonetheless displayed the largest exclusive set of any cell type (∼38,000 transcripts), pointing to a substantial hepatic-specific TE signature absent from all other profiled cell types. Within the neural lineage, NPCs and neurons shared 25,286 transcripts, consistent with a TE program progressively established during neural differentiation. Additionally, neurons also display a distinct group of exclusive transcripts, supporting the literature regarding the prominent role of retroelements in brain tissues (Quinlan and Hall 2010; Evans and Erwin 2021). Notably, iPSCs displayed almost no exclusive transcripts, indicating that the pluripotent TE program is largely a shared subset of what is expressed across differentiated states rather than a unique transcriptional signature. Together, these data demonstrate that TE transcript sharing is lineage-constrained, with cross-cell-type overlap greatest among functionally related cell types, while biologically distinct lineages maintain unique TE transcriptional identities.

**Figure 3.**
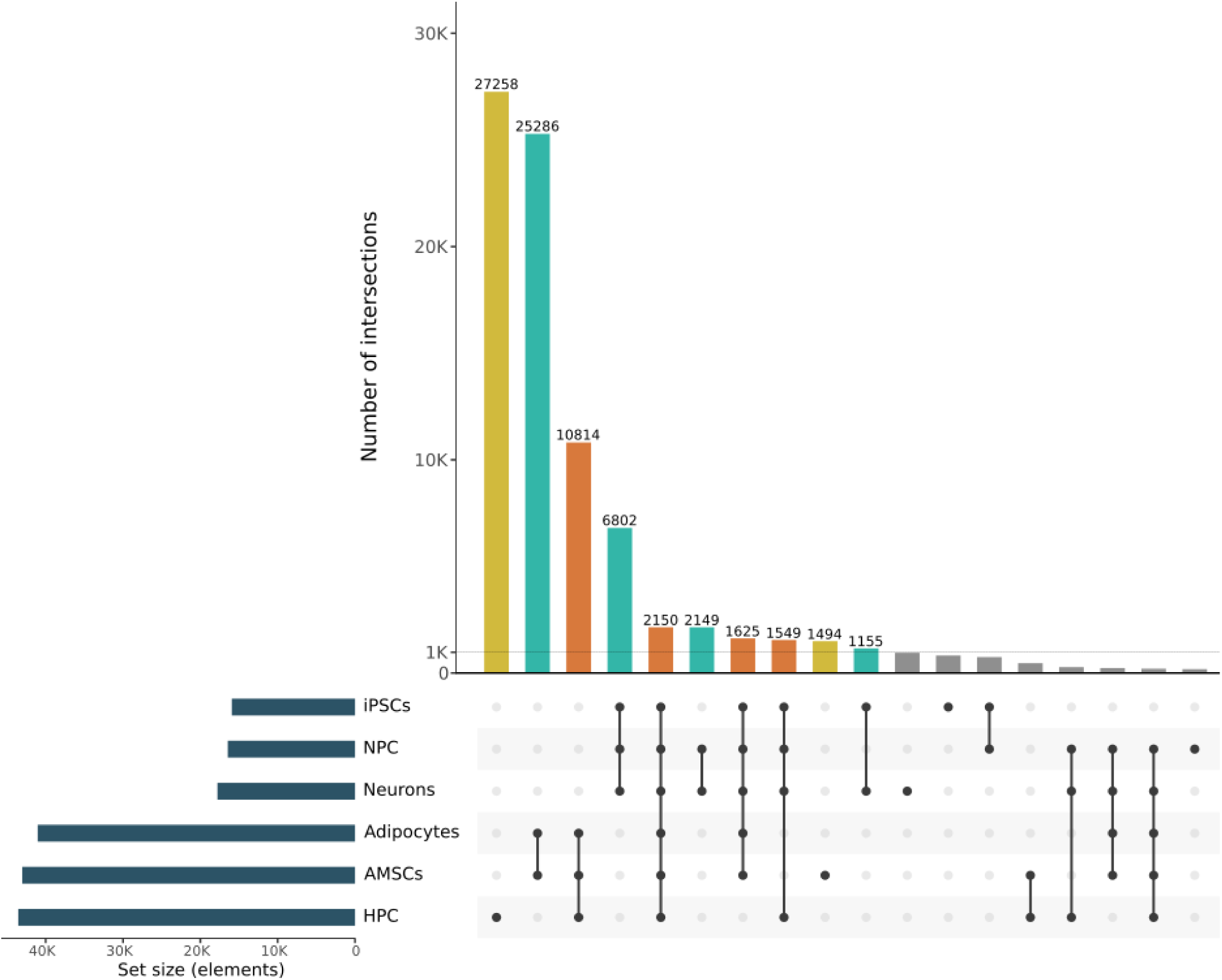
Intersection of transcripts detected by samples. An upset plot representing the number of intersections of transcripts detected across different sample types: induced pluripotent stem cells (iPSCs), neural progenitor cells (NPCs), neurons, adipocytes, adipose-derived mesenchymal stem cells (AMSCs), and hepatic progenitor cells (HPCs). The orange bars indicate the number of intersections for the clusters shown by the connection of the points. The blue bars represent the number of transcripts detected for each cell type.

By examining the sequence and structure of TE-derived transcripts, we identified both the canonical single-exon TE transcripts and more complex isoforms, such as those encompassing multiple TE sequences, and even elements from distinct subfamilies (**Figure 4a**). Moreover, single-TE transcripts were predominantly derived from primate-specific L1 families (**Figure 4b**), especially within neural samples. This highlights the contribution of recently evolved L1 elements as major drivers of TE-derived transcription in these samples (Garza et al. 2023).

**Figure 4.**
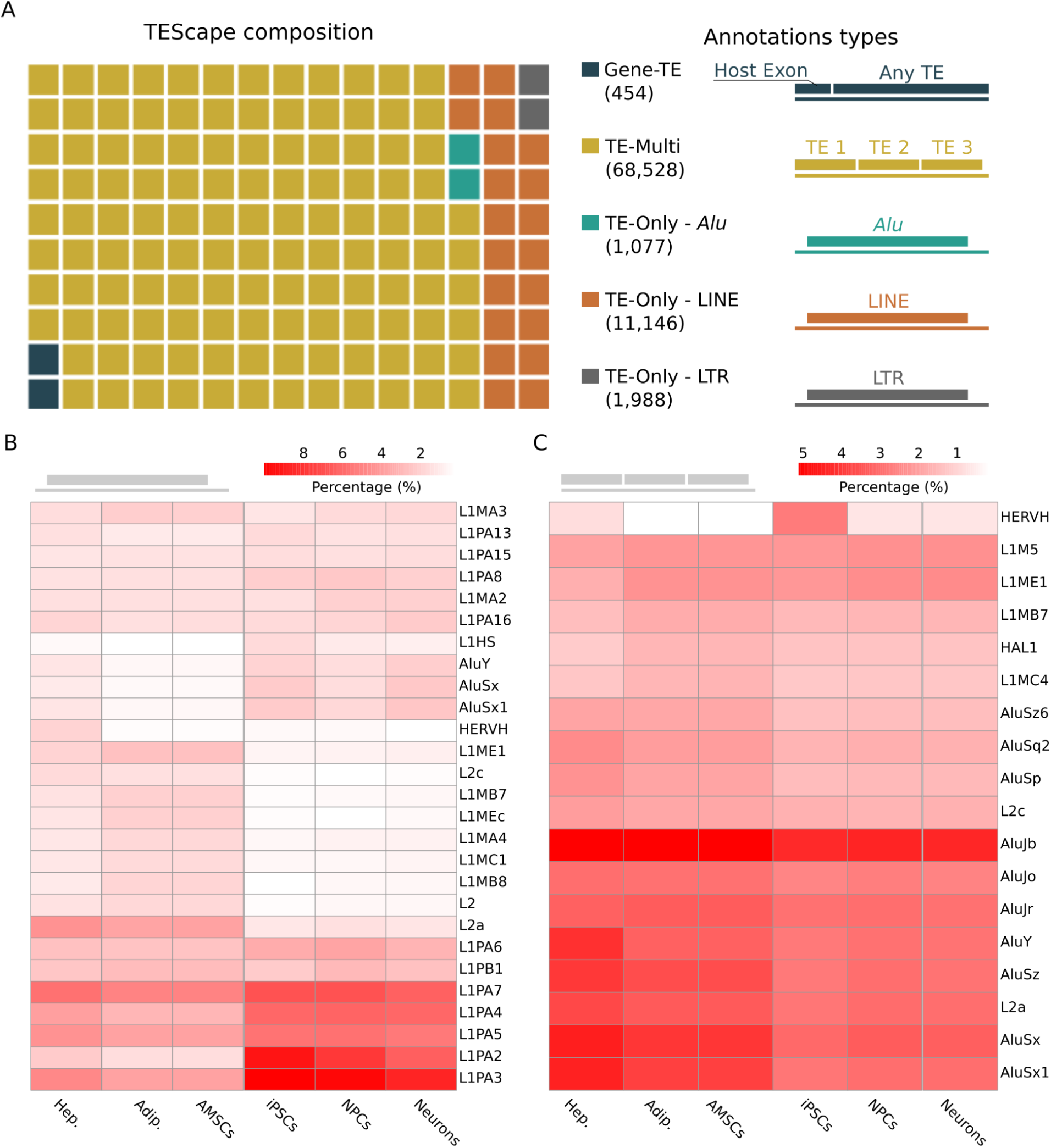
Transposable elements-derived transcripts composition. **A)** Composition of transposable element (TE) classes within the TEscape index. Colors represent distinct annotation types, including gene-overlapping TEs (Gene–TE), multi-element in the same transcript (TE-Multi), and TE-only categories corresponding to *Alu*, LINE, and LTR elements. **B)** Heatmap showing the relative abundance of TE-Only elements (independent of *Alu*, LINE, or LTR classification) across six cell types. **C)** Heatmap of TE-Multi elements, representing loci composed of multiple overlapping TE insertions. Color intensity in panels (B) and (C) indicates families that have at least 1% representation in the transcripts of any sample type.

In contrast to the neural-specific single-TE transcripts, multi-TE transcripts are predominantly composed of *Alu* elements and are primarily detected in non-neural cell types (**Figure 4c**). This pattern reflects the genomic architecture of SINEs, which tend to cluster closely or even overlap within the genome, facilitating their co-transcription. In addition, we observed that the most prevalent mobile elements were *Alu* elements, particularly *AluJb* and *AluJo*. These SINEs are inactive (no longer capable of retrotransposition) (Bennett et al. 2008), and their presence in shared transcripts suggests a secondary function of these elements in the transcription of mobile elements. Additionally, LTR elements (such as LTR7 and HERVH-int) appeared to have greater specificity in iPSCs, potentially linked to the strong promoter present in these elements, which could serve as a transcriptional initiator for adjacent sequences (Babaian and Mager 2016).

In addition, we observed an important evolutionary pattern in these results. Most TEs that contribute to single-TE transcripts are predominantly evolutionarily young elements, such as primate– or human-specific LINE-1 subfamilies (L1P* and L1HS), as well as *AluY* (Tubio et al. 2014; Deininger 2011). In contrast, multi-TE exon transcripts are mainly composed of older elements, some of which are considered transcriptionally inactive in isolation, including mammal-specific LINE-1 subfamilies and LINE-2 elements, in addition to older *Alu* subfamilies such as *AluJ* (Sela et al. 2007)(Sela et al. 2007; Krull et al. 2005).

### TEscape has the potential to improve the interpretation of short-read RNA-Seq data

To further investigate TEscape potential, we performed an expression analysis of short-read RNA-Seq data using 50 randomly selected Liver Hepatocellular Carcinoma (LIHC) tumor samples and 50 matched normal liver tissue samples from The Cancer Genome Atlas (TCGA). We identified transcripts exhibiting canonical alternative splicing (AS) events, resulting in multi-exonic TE-derived isoforms, including cases where TE sequences themselves serve as intronic regions (**Figure 5a**). These spliced transcripts are particularly significant, as the presence of canonical splice junctions enables the recruitment of splicing-dependent export machinery (Reed and Hurt 2002). In contrast, transcripts lacking such splicing signals may be retained in the nucleus due to inefficient export, potentially delaying or impairing their cytoplasmic localization and downstream functional consequences (Valencia et al. 2008).

**Figure 5.**
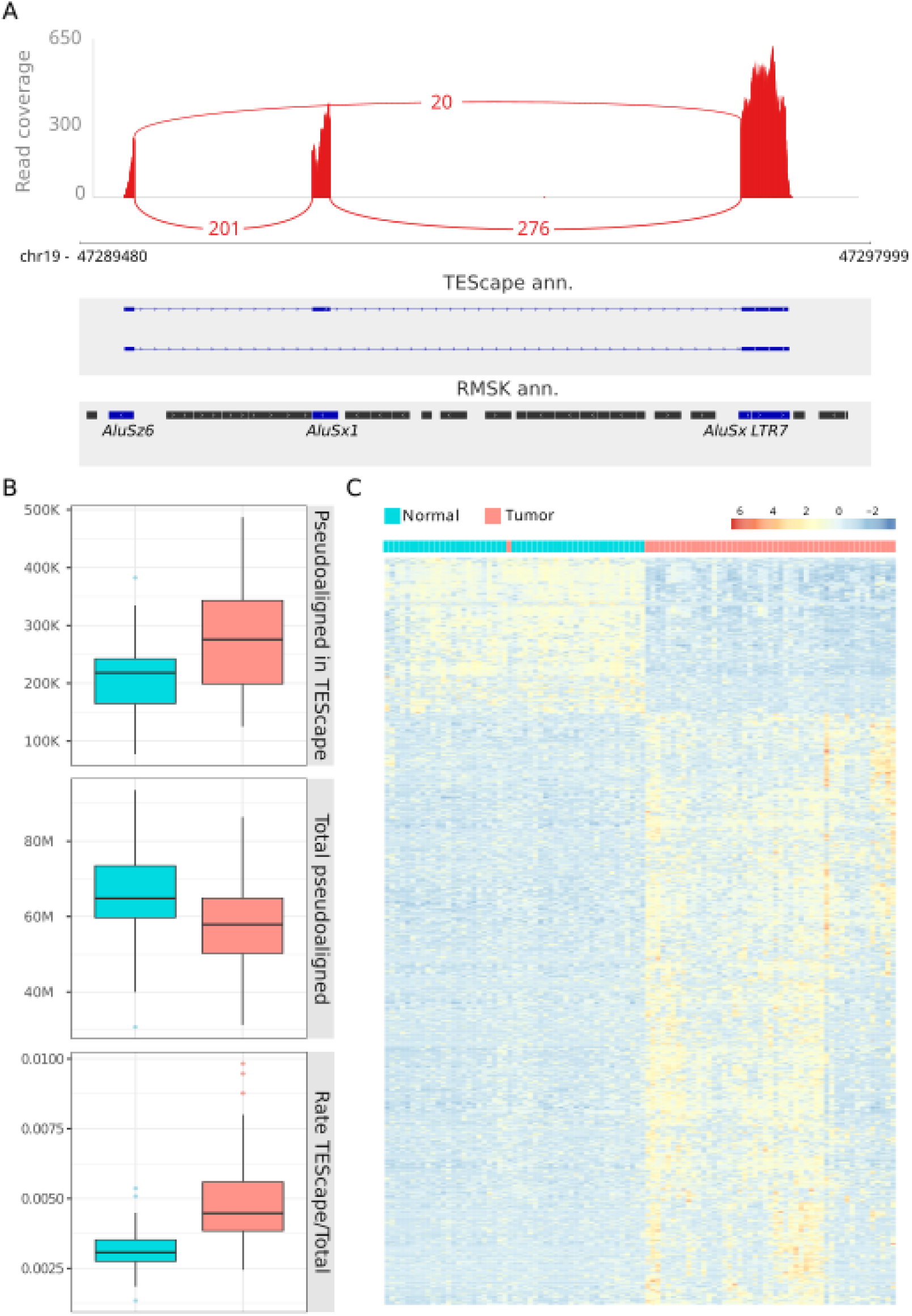
Differential TE-associated transcript activity between normal and tumor samples. **A)** Example of a TE-derived transcript locus identified by TEscape. The upper panel shows read coverage across the region in RNA-seq data, highlighting exons and splice junctions. Middle panels display the corresponding TEscape transcript models detected in hepatocyte (Hep) and neuronal (Neu) datasets, while the bottom panel shows RepeatMasker (RMSK) annotations for individual transposable elements (*AluSx6*, *AluSx1*, *AluSx*, and *LTR7*). **B)** Boxplots compare (from left to right) the number of reads pseudoaligned to TEscape transcripts, the total number of pseudoaligned reads, and the proportion of TEscape reads relative to total reads between normal and tumor samples. **C)** Heatmap showing normalized expression (Z-score) of TEscape transcripts across samples, with distinct clustering separating normal (blue) and tumor (red) groups.

In addition, an apparent increase in TEscape-aligned reads was observed in tumor samples when compared to normal tissues and in the proportion of TEscape reads relative to the overall pseudoaligned reads (**Figure 5b**). This pattern indicates an enhanced representation of TE–derived transcripts in the tumor transcriptome, likely reflecting the transcriptional reactivation of usually silenced repetitive regions or the formation of TE-driven chimeric isoforms (Mercuri et al. 2024). These findings emphasize the relevance of including TE-associated sequences in transcriptome annotations and analyses, as standard RNA-Seq references often fail to capture such noncanonical transcriptional events.

Furthermore, the differential expression analysis revealed 952 upregulated and 232 downregulated TE-derived transcripts. Notably, these transcripts demonstrated a strong capacity to distinguish tumor and normal conditions, as evidenced by the clustering patterns (**Figure 5c**). We identified 1,105 previously unannotated (by Gencode or RefSeq) differentially expressed transcripts recovered by TEscape. This further displays the complexity and potential of TE-derived transcripts. TE-derived transcripts show higher expression levels in tumor samples, which may be related to increased RNA polymerase activity (Nesta et al. 2025; Kong et al. 2019; Johnson et al. 2008).

### Single-cell RNA-Seq data are likewise enhanced by TEscape annotation

Additionally, we conducted a 10x 3’ short read scRNA-Seq analysis using the same library as the MAS-ISO-Seq experiment on hepatic cell samples. In this analysis, we used as a mask the classifications assigned to the cells based on protein-coding signatures from a previous analysis.

We used the previous cell classification to confirm that the expression of MEs alone allows for the identification of distinct expression profiles by cell type and we were also able to identify the expression of several TE-derived transcripts that are directly associated with specific cell types. (**Figure 6a**). It is important to highlight that, although many of the transcripts detected in the MAS-ISO-Seq data are shared across multiple cells, the expression quantification for these cases is likely distinct, leading to the creation of different expression patterns.

**Figure 6.**
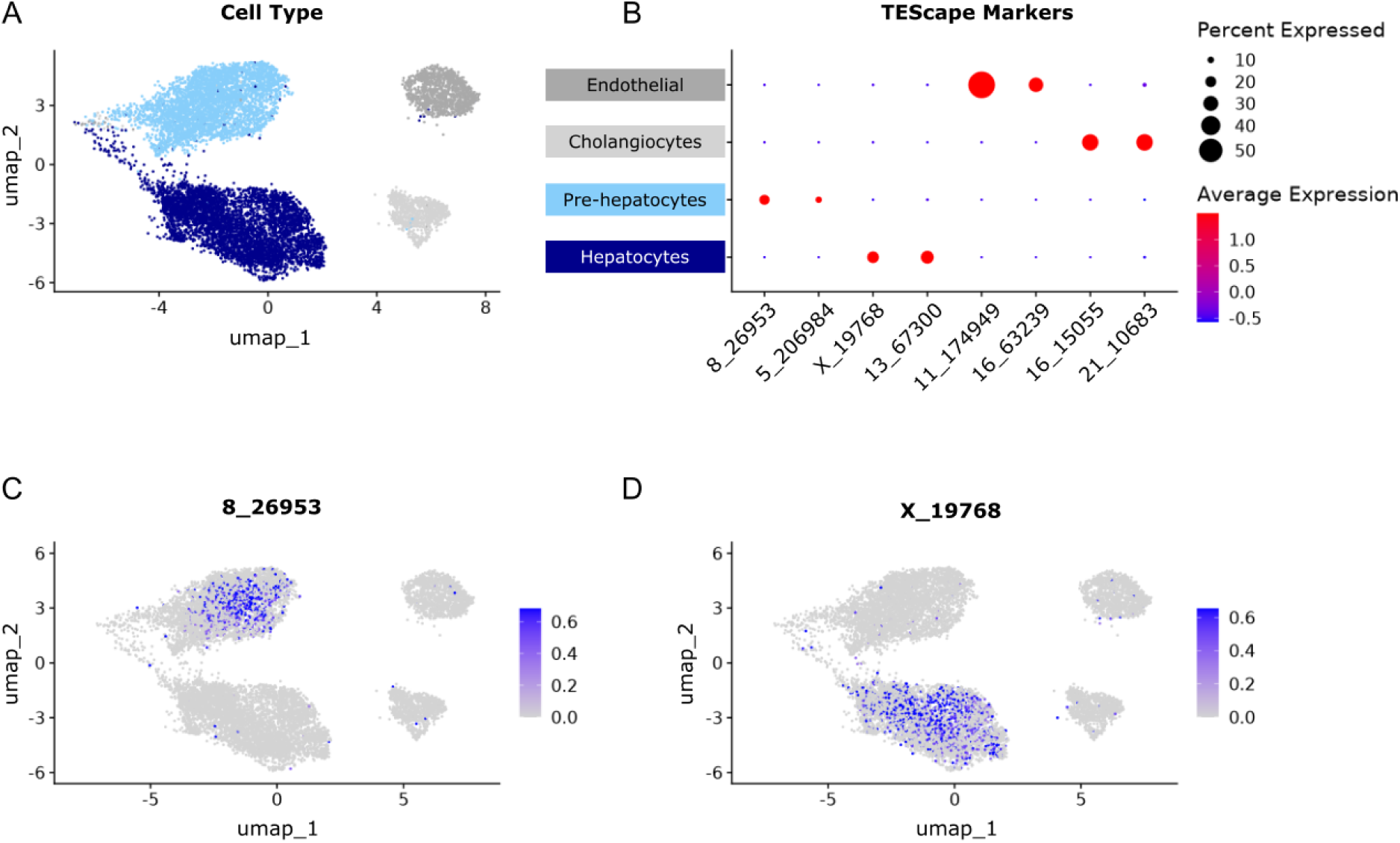
Mobile Element Expression Profiles by Cell Type in Hepatic Samples. **A)** UMAP visualization of 10X scRNA-Seq data from hepatic progenitor cells based on the expression of mobile elements (MEs) in differentiated liver cells. Hepatocytes are shown in dark blue, pre-hepatocytes in light blue, cholangiocytes in light gray, and endothelial cells in gray. **B)** Dot plot showing the main TE-derived transcript markers identified by TEscape across cell populations. Dot size represents the percentage of cells expressing each transcript, while color intensity indicates the average expression level. **C–D)** Spatial distribution of the representative TE-derived transcripts *8_26953* and *X_19768* across the UMAP embedding. Expression of *8_26953* is enriched in pre-hepatocytes, whereas *X_19768* shows higher expression in hepatocytes, highlighting cell type-specific expression patterns.

Next, we evaluated the expression profiles of TE-derived transcripts identified by TEscape across these cell populations. Differential expression analysis revealed distinct TE-derived transcript markers associated with specific liver cell types (**Figure 6b**). In particular, transcripts *8_26953* (TE-Multi: L1PA12|L1M5|L1M5|L2c) and *5_206984* (TE-Multi: L1PA5|*AluSq2*) showed enriched expression in pre-hepatocytes, whereas *X_19768* (TE-Multi: *AluJo*|MIR|L1PA8) and *13_67300* (TE-Multi: L1PA14|L1PA15) were predominantly expressed in hepatocytes. In contrast, *16_15055* (TE-Multi: MamGypLTR1c|L1MEb|*AluSq*|L1MEb|L1MEb|MIRb) and *21_10683* (TE-Only: HERVH-int) displayed preferential expression in cholangiocytes, while *11_174949* (TE-Multi: *AluSx*|L1ME4b) and *16_63239* (TE-Multi: L1M6|*AluJr*) were mainly associated with endothelial cells. These results suggest that TE-derived transcripts are not uniformly expressed across liver populations, but instead exhibit strong cellular specificity.

To further characterize these patterns, we visualized the spatial distribution of representative TE-derived transcripts across the UMAP embedding. Transcript *8_26953* showed restricted expression predominantly within the pre-hepatocyte cluster (**Figure 6c**), while *X_19768* was primarily detected in hepatocytes (**Figure 6d**). The localized expression of these transcripts supports the idea that TE-derived transcriptional activity reflects cellular identity and may contribute to cell type-specific regulatory programs.

Overall, these findings demonstrate that TEscape is able to identify TE-derived transcripts with highly specific expression patterns at single-cell resolution, revealing potential biomarkers associated with distinct liver cell populations.

## DISCUSSION

The reference status of genomes is defined by comprehensive annotations, such as those provided by the GENCODE and RefSeq projects, which build upon large-scale efforts like ENCODE to map the complexity of eukaryotic transcription. Yet current reference annotations, such as GENCODE v47 (Mudge et al. 2025), have captured only a fraction of the transcript diversity driven by TEs. The identification of more than 83,000 TE-derived transcripts by TEscape, many of which exhibit substantial structural novelty, dramatically expands the known repertoire of the human transcriptome, a point underscored by the limited overlap with existing annotations, where only 1,258 transcripts (773 genes) from the TEscape catalog were previously annotated in GENCODE. Together, these findings indicate that the vast majority of TE-derived transcripts represent an underrepresented expansion of the human transcriptional landscape, underscoring the need to reassess the boundaries of TE reference transcriptome annotation.

This observation echoes findings from the ENCODE project, which has consistently shown that existing catalogs capture only a subset of possible combinations of TSSs, exon junction chains (ECs), and TTSs, for instance, ENCODE LR-RNA-seq identified 76,469 transcripts with exon junctions absent from GENCODE v29 and v40. TEscape adds a new dimension to this complexity by revealing multi-exon transcripts (up to 16 exons) and multi-TE transcripts, a level of structural diversity that canonical annotations have historically underrepresented, in part due to their emphasis on a single principal transcript per gene, as in MANE Select or RefSeq Select sets (Pozo et al. 2022), and the technical limitations of short-read RNA-seq in reconstructing full transcript structures (Byrne et al. 2019; David et al. 2022). The ability of TEscape to robustly characterize this diversity is inherently tied to long-read RNA sequencing, here implemented via MAS-ISO-seq, which enables reconstruction of complete transcript structures and mirrors the approach adopted by ENCODE in its ENCODE4 phase through a triplet-based TSS–EC–TES framework; the identification of 18.0% of TSSs, 37.3% of ECs, and 22.1% of TESs as novel features by ENCODE (Reese et al. 2023), together with the TE-derived transcripts uncovered here, reinforces the view that transcript diversity is an intrinsic and evolvable property of genes that must be systematically cataloged. Consistent with this, TEscape refined the resolution of transcription start and end sites, demonstrating that TEs can initiate or terminate transcription beyond their own boundaries, producing “leaky expression”, an approach analogous to GENCODE’s use of Capture Long-read Sequencing (CLS3), which achieved a three– to six-fold increase in annotated lncRNA transcripts (Kaur et al. 2024), collectively underscoring the effectiveness of LR-RNA-seq in capturing complex and noncoding RNA classes with high accuracy.

The role of TE-derived transcripts is not merely structural but also regulatory: over 85% harbor regulatory expression markers, including promoters, proximal and distal enhancers, and CTCF-binding sites, as identified using ENCODE datasets and TSS databases, suggesting broad functional relevance for these newly cataloged elements. This integration of regulatory features reflects the future trajectory of large-scale annotation projects; RefSeq has acknowledged the importance of non-genic elements through the RefSeq Functional Elements initiative (Farrell et al. 2022; Goldfarb et al. 2025), while GENCODE is advancing the concept of the “extended gene” by refining UTR annotations and introducing a promoter window (1,000 bp upstream of the MANE Select TSS) to link gene annotations with experimental regulatory data from ENCODE, a framework into which the high frequency of regulatory markers in TEscape transcripts directly argues for their incorporation. The clinical implications of excluding such noncanonical transcripts are equally substantial (Zhang et al. 2020; Guardia et al. 2025): preliminary analysis of hepatocellular carcinoma (LIHC) revealed that 1,105 transcripts absent from both GENCODE and RefSeq, yet recovered by TEscape, were differentially expressed and capable of distinguishing tumor from normal tissue, directly challenging the prevailing clinical practice of restricting analysis to a single representative transcript per gene (e.g., MANE Select), a constraint that risks obscuring critical transcript-level distinctions and confounding the interpretation of variant effects, while demonstrating that incorporation of the TEscape index into short-read RNA-seq quantification offers an immediate path to improving the interpretation of existing transcriptomic datasets.

In conclusion, the TEscape catalog constitutes a transcript-level reference atlas that fundamentally redefines the boundaries of the annotated human transcriptome, revealing transposable elements as pervasive contributors to transcriptional and regulatory complexity rather than genomic noise. The structural novelty, regulatory enrichment, and clinical relevance of the more than 83,000 TE-derived transcripts cataloged here collectively make the case that canonical reference annotations, however comprehensive, remain insufficient representations of the transcriptional landscape. Integrating TEscape into existing reference frameworks is therefore not merely an incremental refinement but a necessary step toward accurate quantification, variant interpretation, and disease-relevant transcriptomic analysis across diverse cellular contexts. As long-read sequencing technologies continue to mature and large-scale initiatives such as ENCODE, GENCODE, and RefSeq expand their regulatory and isoform-level scope, TE-derived transcripts must be recognized as first-class constituents of the human transcriptome, a reclassification with broad implications for genomics, clinical diagnostics, and our understanding of the human genome.

## METHODOLOGY

### Preprocessing MAS-ISO-seq datasets

The MAS-ISO-Seq data were preprocessed using the Pigeon (version 4.1.0) pipeline (https://isoseq.how/), which was originally developed for ISO-Seq data but has since been updated to accommodate the specific requirements of MAS-ISO-Seq. The preprocessing workflow began with the demultiplexing of degenerate primers using the ––isoseq parameter and the 10x 3′ kit primer sequences. For libraries derived from single-cell protocols, unique molecular identifiers (UMIs) and cell barcodes were assigned using the ––design T-12U-16B parameter during the tagging step. These single-cell libraries also underwent barcode correction and duplicate removal to ensure accurate quantification and cell-level resolution.

Subsequently, read correction was performed in the refine step, with amplification primers provided as input to improve base-level accuracy. Following these preprocessing steps, the reads were aligned to the human reference genome (hg38) using the pbmm2 aligner (version 1.16.0) (https://github.com/PacificBiosciences/pbmm2) configured with the ––preset ISOSEQ and ––sort parameters to optimize for full-length transcript alignment.

### Annotation

To annotate the retroelement transcriptome, we first assembled the general transcriptome of each sample using the IsoQuant (version 3.6.1) tool (Prjibelski et al., 2023), employing Gencode v47 (Mudge et al., 2025) as the reference transcriptome. The assembly was performed with the following parameters: ––complete_genedb, ––data_type pacbio_ccs, and ––report_novel_unspliced true. Following transcriptome assembly, we applied stringent filtering using SQANTI3 (version 5.2.2) (Pardo-Palacios et al. 2024) to eliminate transcripts with features indicative of technical artifacts. Specifically, isoforms were flagged as potential intrapriming artifacts if the genomic region within 20 bp downstream of the annotated transcription termination site (TTS) contained ≥12 adenines, corresponding to a stretch with ≥60% adenine content. Additionally, isoforms were retained only if they lacked reverse transcription switching (RT-switching) junctions and contained exclusively canonical splice junctions.

After filtering, we identified high-confidence transcripts (minimally represented with 5 reads or more) and assessed their origin relative to mobile genetic elements. To do this, we developed custom in-house scripts to compare transcript coordinates against annotated genomic locations of retroelements, including *Alu*, LINE1, SVA, and LTR families, as well as retrocopies. Annotation data for these elements were sourced from RepeatMasker (http://www.repeatmasker.org). A transcript was classified as TE-overlapping transcript if it overlapped ≥60% with any annotated retroelement or retrocopy region. To ensure robustness, we applied additional thresholds: each transcript had to be supported by at least five reads, and for samples derived from the Village dataset, transcripts were required to be present in at least two donors.

### TEscape reference building

To construct a comprehensive reference transcriptome, we integrated the high-confidence transcripts identified through our analytical pipeline into the existing Gencode v47 annotation. This merged dataset was compiled into a custom GTF file, which serves as an enhanced reference for downstream transcriptomic analyses.

### Bulk RNA-Sequencing quantification

We applied the newly generated reference transcriptome using Kallisto (version 0.48.0) (default parameters) (Sullivan et al. 2025) as the quantification tool for short-read data in two distinct conditions. The first condition involved 50 random RNA-Seq samples of hepatocellular carcinoma (LIHC) and 50 normal liver tissue samples from The Cancer Genome Atlas (TCGA). For differential expression analysis (LFC >= 2 and *p-adj* <= 0.05) and PCA, we used the DESeq2 package (version 1.38.3) (Love et al. 2014).

## 10X data quantification

For scRNA-Seq data, we used the CellRanger tool (version 8.0.1, default parameters) as the quantifier with paired data from hepatic progenitor cells, for this step, we excluded cells assigned to more than one donor during demultiplexing. For the analysis of this quantification, we employed the Seurat package (version 5.3.0) (Love, Huber, and Anders 2014; Hao et al. 2024). With the matrix generated by the 10x Genomics Cell Ranger (Zheng et al. 2017), we performed the following steps: (1) Removed cells with high levels of mitochondrial gene expression (> 5%), low numbers of expressed mobile elements (< 5%), and low n_RNAFeatures (< 200). (2) Normalized the matrix with all genes. (3) Clustered the cells based solely on the expression of mobile elements, using the 5000 most variable mobile elements and 50 principal components in UMAP dimensionality reduction. For this dataset, we used a prior annotation of cell types identified by protein-coding genes to attempt clustering of cell types based solely on the expression of mobile elements.

### AMSCs cell culture, differentiation and nuclei isolation

Subcutaneous adipose tissue was collected from 2 donors undergoing liposuction (donor 1, age: 33, sex: female and BMI: 26.8; Donor 2, age: 40, sex: female and BMI: 35.6). Each donor provided written informed consent before adipose tissue biopsies were collected. Primary AMSCs were isolated, cultured and differentiated as previously described (PMID: 37492099, PMID: 22057455). Briefly, cells were maintained in proliferation media (PM: DMEM/F-12 supplemented with 1% Penicillin-Streptomycin, 33 μM Biotin, 17 μM Pantothenate, 0.13 μM Insulin, 10 ng/ml EGF, 1 ng/ml FGF and 2.5% FBS) and differentiation was initiated in fully confluent cells by changing PM to induction media (IM: DMEM/F-12 supplemented with 1% Penicillin-Streptomycin, 33 μM Biotin, 17 μM Pantothenate, 0.86 μM Insulin, 1 nM T3, 0.1 μM Hydrocortisone, 0.01 mg/ml Transferrin, 0.17 % fatty acid free BSA, 2% FBS, 1 μM Rosiglitazone, 25 nM Dexamethasone and 0.25 mM IBMX). Following 3 days, IM was changed to differentiation media (DM: DMEM/F-12 supplemented with 1% Penicillin-Streptomycin, 33 μM Biotin, 17 μM Pantothenate, 0.86 μM Insulin, 1 nM T3, 0.1 μM Hydrocortisone, 0.01 mg/ml Transferrin, 0.17 % fatty acid free BSA and 2% FBS) for a total period of 14 days.

Cells from each donor were cultured in PM independently in a T-75 flask until around 80% confluent, at which point AMSCs from two donors were pooled at equal proportions and seeded at a density of 400,000 cells per well of a 6 well plate. After pooling, fully confluent cells were differentiated as described above and nuclei were isolated at day 0 and 14 of differentiation. First, AMSCs were washed once with pre-warmed PBS and incubated with 2 ml per well of serum-free DMEM/F12 supplemented with 50 μg/ml DNase I (Stem Cell Tech, #07900) for 10 min at 37°C in a cell culture incubator. Next, cells were placed on ice, washed twice with ice-cold PBS and collected in 500 µl of ice-cold Nuclei Lysis Buffer (5 mM CaCl2, 3 mM Mg(Ac)2, 10 mM Tris pH 7.5, 320 mM Sucrose, 0.1 mM mM EDTA, 0.1% IGEPAL CA-630, 1 mM DTT and 1 U/μl RNase inhibitor) into a pre-chilled 1.5 ml low protein binding tube (Eppendorf, #022431081). Samples were mechanically disrupted by pipetting 15 times, followed by 3 min incubation on ice and subsequent 15 strokes with a tight pestle in a glass Dounce homogenizer on ice. The resulting cell lysate was transferred to a 5 ml tube on ice, ice-cold Wash Buffer (1X PBS, 2% BSA and 1 U/μl RNase inhibitor) was added for a 4 ml final volume and mixed gently by inverting the tube five times. Samples were next centrifuged for 5 min at 500 rcf at 4°C in a swinging bucket centrifuge with the acceleration and brake set at 50% of the maximal setting. The supernatant was discarded, the nuclei pellet was resuspended in 2 ml of Freezing Buffer (Wash Buffer supplemented with 10% DMSO) and 1 ml aliquots transferred to a cryovial for overnight freezing in a freezing container at -80°C. Samples were thawed in a room temperature water bath and filtered through a 30 μm cell strainer (Sysmex America, #04-004-2326) into a pre-chilled 1.5 ml low protein binding tube on ice. Nuclei were permeabilized with 0.5% Tween-20 and 0.4% Digitonin, followed by pipette mixing five times and incubation on ice for 10 min. Next, nuclei were washed by centrifugation for 5 min at 500 rcf at 4°C in a swinging bucket centrifuge with the acceleration and brake set at 50% of the maximal setting. The supernatant was discarded, the nuclei pellet was resuspended in 150 μl of 1X Nuclei Buffer (10x Genomics, PN-2000207), visually inspected and counted in an hemocytometer using trypan blue staining. Samples were then centrifuged for 5 min at 500 rcf at 4°C in a swinging bucket centrifuge with the acceleration and brake set at 50% of the maximal setting, supernatant was discarded and nuclei pellet was resuspended in appropriate volume of 1X Nuclei Buffer to immediately proceed with the Chromium Next GEM Single Cell Multiome ATAC + Gene Expression kit (10x Genomics, PN-1000283; protocol version CG000338 Rev F). MAS-ISO-seq was next performed using cDNA samples from snRNA-seq libraries.

## Acknowledgment

P.A.F.G. acknowledges support from the São Paulo Research Foundation (FAPESP; grants #2018/15579-8 and #2025/18246-3). R.L.V.M. acknowledges support from FAPESP (fellowship #2020/02413-4 and #2023/11391-2). D.M.M. acknowledges support from FAPESP (fellowship #2025/11174-7). R.L.V.M. were supported by fellowship from the Young Scientist program, Hospital Sírio-Libanês. It was also partially supported by funds from CNPq, Serrapilheira Foundation, and Hospital Sírio-Libanês to PAFG. RLVM would also like to express the deepest gratitude to his parents, Luiz M. P. Mercuri and Marly M. R. Vieira, whose constant love, encouragement, and emotional support carried him through every stage of this work. Though they may no longer be here to witness its completion, he believes that, wherever they are, they would be happy to see this journey reach its conclusion.

