## Supplementary Material for "TEscape: Defining the human transposable element transcriptome using multiplatform long-read sequencing"

**Supplementary Table 1. Reads count by sample in MAS-ISO-Seq data.**

| Cell type | Sample type | Experiment (MAS-ISO-Seq) | # of Reads |
|---|---|---|---|
| Hepatic Progenitor | Village | scRNA-Seq | 58,620,946 |
| AMSCs (Day0) | 2 donors | snRNA-Seq | 45,032,127 |
| Adipocytes (Day14) | 2 donors | snRNA-Seq | 50,046,268 |
| iPSCs | iPSCs | Bulk RNA-Seq | 26,378,095 |
| NPCs | iPSCs | Bulk RNA-Seq | 24,801,689 |
| Neurons | iPSCs | Bulk RNA-Seq | 30,717,156 |
| TOTAL | | | 235,596,281 |

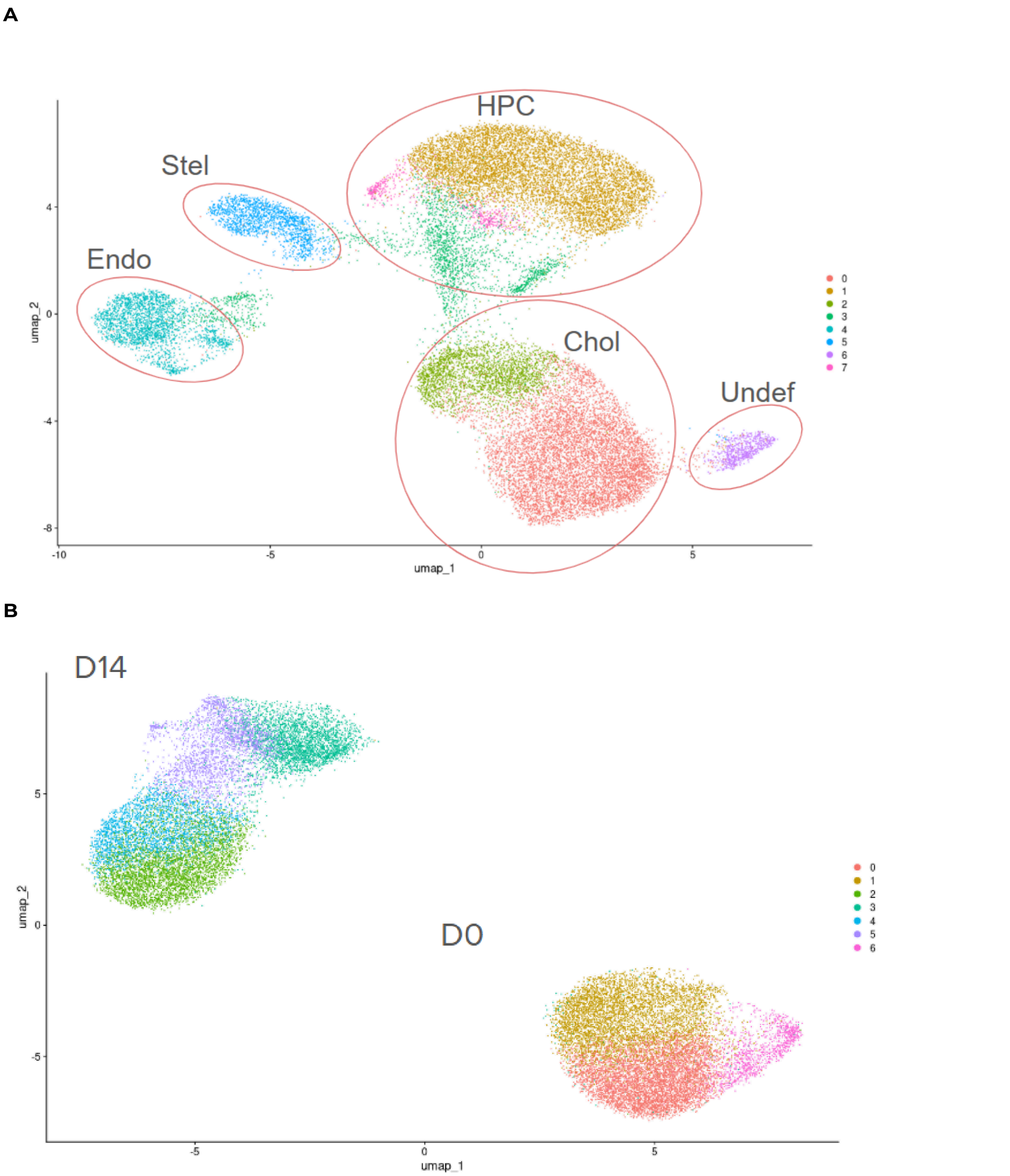


**Supplementary Figure 1. UMAP visualization of gene expression across cell types and conditions. A)** UMAP embedding showing the cell populations identified based on gene profiles (iPSCs - scRNA-Seq), including hepatocyte progenitor cells (HPC), cholangiocytes (Chol), endothelial cells (Endo), stellate cells (Stel), and an undefined cluster (Undef). Clusters are highlighted and annotated according to their gene identity. **B)** UMAP projection by experimental condition (AMSCs - snRNA-Seq), illustrating the separation between Day 0 (D0) and Day 14 (D14), indicating temporal shifts in gene expression patterns.

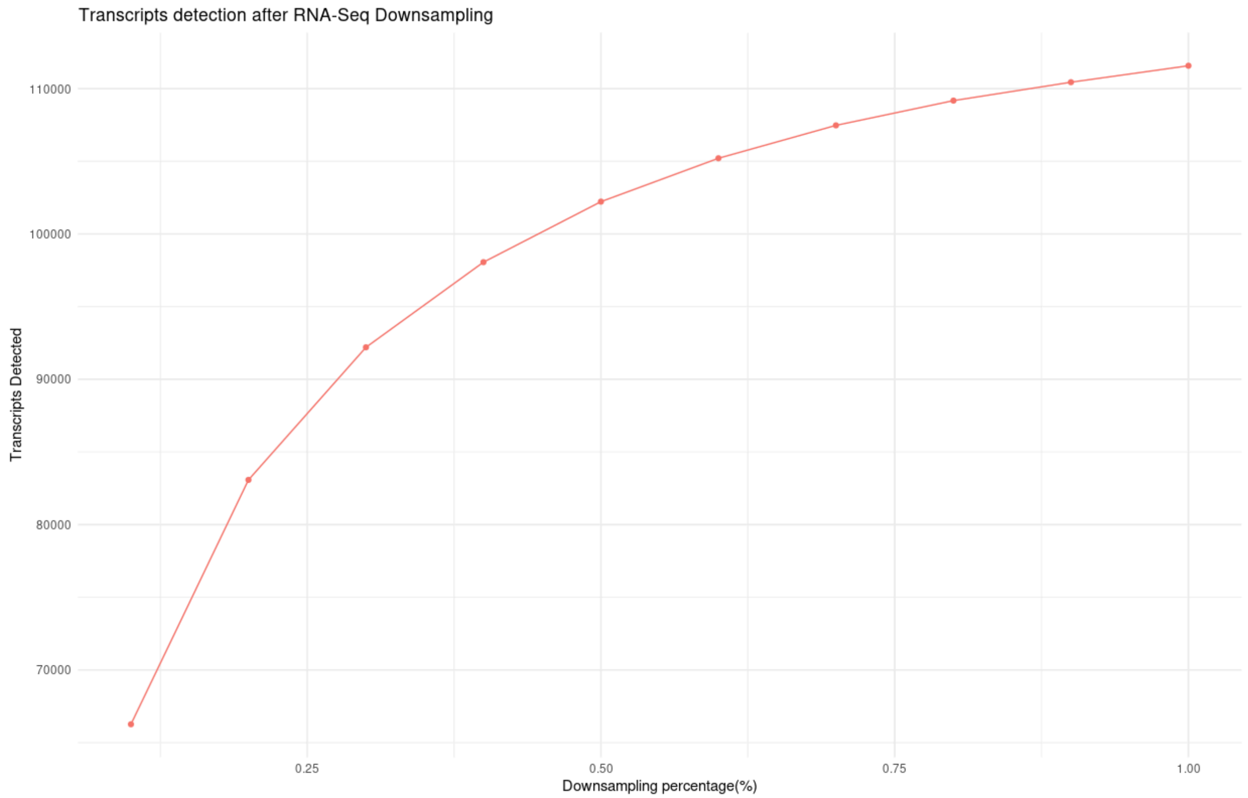


**Supplementary Figure 2. Downsampling analysis that our sequencing libraries achieved approximately 90% of transcript detection limits across all samples.**

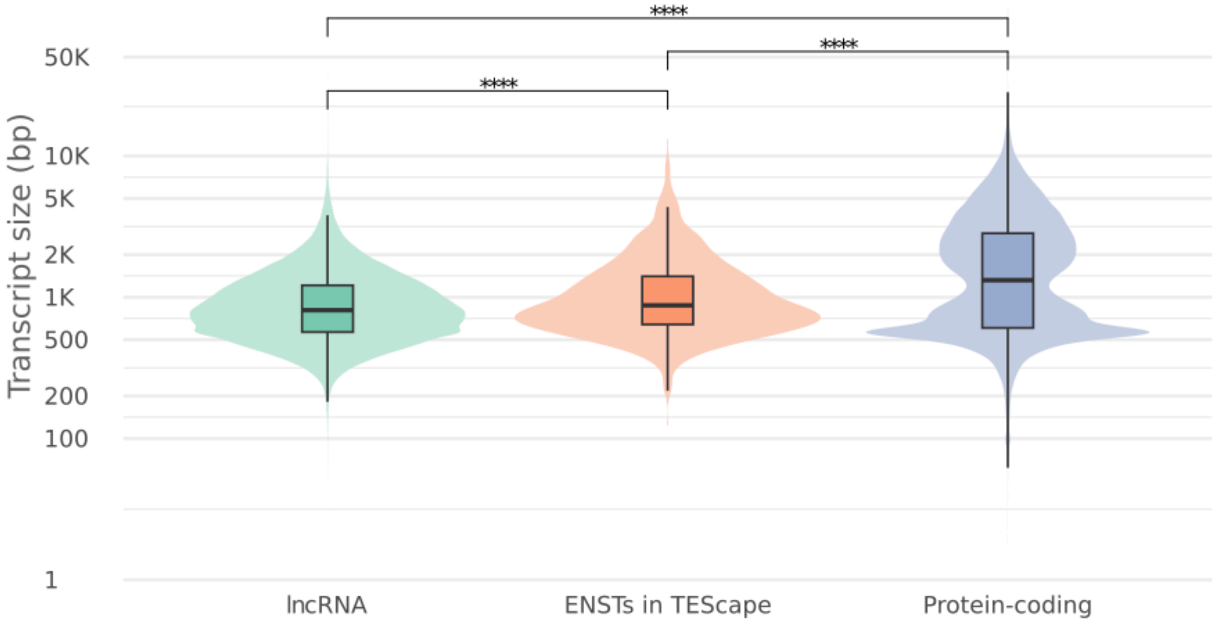


**Supplementary Figure 3. Gencode v47 transcripts comparison between Gencode transcripts.**

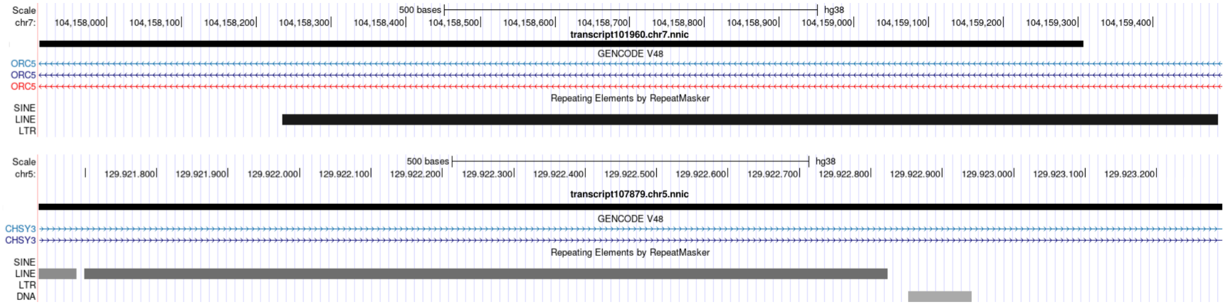


**Supplementary Figure 4. Examples.**
